# mAIcrobe: an open-source framework for high-throughput bacterial image analysis

**DOI:** 10.1101/2025.10.21.683709

**Authors:** António D. Brito, Dominik Alwardt, Beatriz de P. Mariz, Sérgio R. Filipe, Mariana G. Pinho, Bruno M. Saraiva, Ricardo Henriques

## Abstract

Quantitative analysis in bacterial microscopy is often hindered by diverse cell morphologies, population heterogeneity, and the requirement for specialised computational expertise. To address these challenges, mAIcrobe is introduced as an open-source framework that broadens access to advanced bacterial image analysis by integrating a suite of deep learning models. mAIcrobe incorporates multiple segmentation algorithms, including StarDist, CellPose, and U-Net, alongside comprehensive morphological profiling and an adaptable neural network classifier, all within the napari ecosystem. This unified platform enables the analysis of a wide range of bacterial species, from spherical *Staphylococcus aureus* to rod-shaped *Escherichia coli*, across various microscopy modalities within a single environment. The biological utility of mAIcrobe is demonstrated through its application to antibiotic phenotyping in *E. coli* and the identification of cell cycle defects in *S. aureus* DnaA mutants. The modular design, supported by Jupyter notebooks, facilitates custom model development and extends AI-driven image analysis capabilities to the broader microbiology community. Building upon the foundation established by eHooke, mAIcrobe represents a substantial advancement in automated and reproducible bacterial microscopy.

## Introduction

Microscopy remains fundamental to microbial cell biology; however, quantitative analysis of bacterial images presents significant challenges. These include population heterogeneity, the small size of bacterial cells, morphological diversity among species, and the range of imaging techniques employed. Manual analysis constitutes a major bottleneck, as it is time-consuming, subjective, and susceptible to human error, thereby limiting research throughput and reproducibility.

To overcome these limitations, we developed mAIcrobe, a comprehensive framework for bacterial image analysis. It supports multiple bacterial species, various microscopy modalities, and flexible, customisable analysis workflows. By integrating various segmentation methods, quantitative morphological measurements, and an adaptable classification model, mAIcrobe provides a powerful tool for a broad range of studies in bacterial cell biology. We have made our work accessible through the napari-mAIcrobe plugin, which is accompanied by Jupyter notebooks (Table S1) to facilitate the training of custom classification models usable within the user-friendly napari ecosystem.

The field has seen several automated image analysis tools tailored for bacterial images, from ImageJ (1) plugins like MicrobeJ (2) to standalone software such as SuperSegger (3) and Oufti (4). Our own contribution, eHooke (5), provided an open-source solution for the semi-automated analysis of cocci, particularly *Staphylococcus aureus*. Although eHooke was a valuable tool for studying the cell cycle in spherical bacteria, it was architecturally constrained, limiting its application to other morphologies and making integration with new deep learning models challenging. DeepBacs (6) has also made available several state-of-the-art artificial neural-network models tailored for bacterial microscopy using the ZeroCostDL4Mic platform (7). The rapid evolution of bioimage analysis, coupled with the broad adoption of the napari ecosystem (8), presented a clear opportunity to engineer a more powerful and extensible framework built on the modern scientific Python stack.

We seized this opportunity to design mAIcrobe (Fig. 1), a next-generation platform prioritising versatility and performance. The design philosophy focused on overcoming morphological constraints, enabling the selection of optimal segmentation algorithms, and facilitating the rapid adaptation of deep learning models to address emerging biological questions. The napari framework was selected as the foundation for mAIcrobe due to its modular architecture and interactive visualisation capabilities, which align with these objectives.

**Fig. 1.**
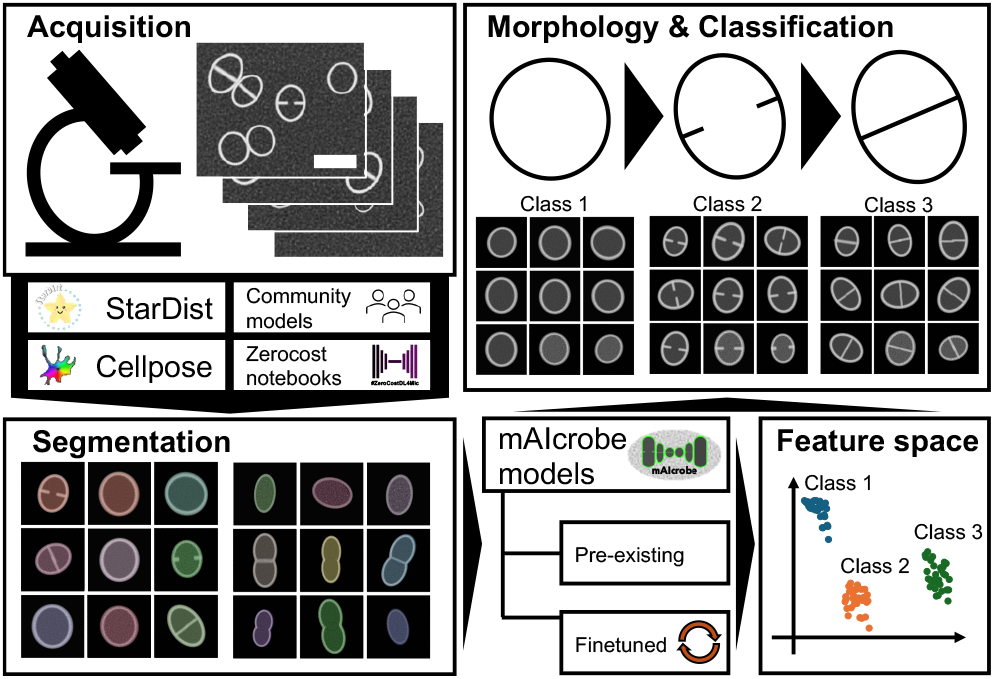
mAIcrobe workflow. After image acquisition, the napari-mAIcrobe plugin facilitates analysis through a user-friendly interface for segmentation, morphological measurements, and classification using a variety of pre-trained or custom models.

## Results

Developed within the napari plugin ecosystem, mAIcrobe provides an intuitive and extensible platform for bacterial image analysis. The framework integrates image segmentation, morphological measurement, and classification into a unified workflow. Its modular design enables the selection of segmentation models and classification strategies tailored to specific experimental requirements (Table S2). A central feature of the napari-mAIcrobe plugin is its support for real-time visualisation and dynamic parameter adjustment, facilitating optimisation of image processing across diverse bacterial species and microscopy setups (Fig. S1). The following sections illustrate these capabilities through selected biological applications.

***Segmentation***. mAIcrobe features a flexible segmentation engine designed to accommodate diverse bacterial species and microscopy modalities (Fig. 2 and Table S3). The framework integrates several leading segmentation approaches, including StarDist (9), CellPose (10), and custom U-Net (11) models trained using the ZeroCostDL4Mic framework (7). This multi-model strategy enables the selection of the most suitable algorithm for specific experimental conditions and bacterial morphologies, thereby ensuring high-quality segmentation. Unlike tools limited to particular morphologies, mAIcrobe supports the analysis of rod-shaped, spherical, and other bacterial forms within a single framework.

**Fig. 2.**
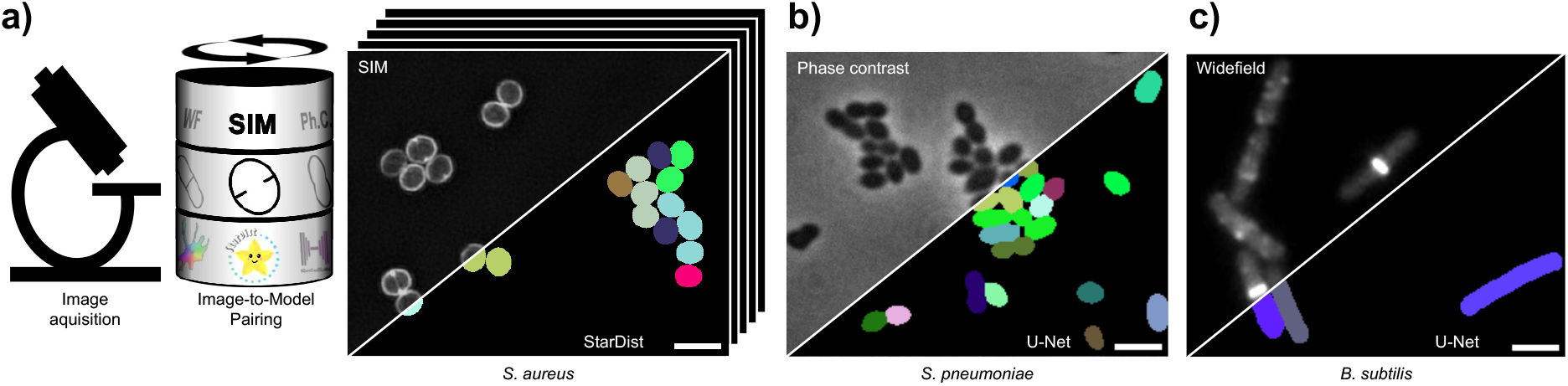
mAIcrobe segmentation capabilities. The platform can perform segmentation for a variety of bacterial species, segmentation models, and microscopy modalities. a) SIM image of *S. aureus* JE2 strain labeled with membrane dye NileRed, with cells segmented using a StarDist model. b) Phase-contrast image of *Streptococcus pneumoniae* segmented with a U-Net trained via ZeroCostDL4Mic. c) Conventional fluorescence widefield microscopy of a *Bacillus subtilis* strain expressing FtsZ-GFP also segmented using a U-Net trained via ZeroCostDL4Mic. All scale bars are 2 µm.

This versatility is demonstrated in several applications. In structured illumination microscopy (SIM) images of *S. aureus* labelled with membrane dye NileRed, the StarDist model achieves accurate cell boundary detection (Fig. 2a). For phase-contrast microscopy of *Streptococcus pneumoniae*, U-Net models trained with ZeroCostDL4Mic provide reliable segmentation of cells (Fig. 2b). The framework also processes conventional widefield fluorescence images, segmenting *Bacillus subtilis* expressing FtsZ-GFP (Fig. 2c) (6). By integrating these models within a single interface, mAIcrobe eliminates the need to switch between software packages, thereby streamlining the identification of optimal segmentation approaches.

### Morphological Measurements

Beyond segmentation, mAIcrobe performs quantitative morphological analysis of bacterial cell properties across diverse experimental conditions (Table S2). From segmented cells, the framework extracts key morphological parameters, including cell area, perimeter, and eccentricity, alongside multi-channel fluorescence intensity measurements. This quantitative data provides a solid basis for characterising cellular responses to drug treatments or genetic modifications. To ensure interoperability and support reproducible research, all results can be readily exported to standard formats, such as CSV, for downstream statistical analysis and visualisation.

A practical application is the detection and characterisation of drug-induced morphological changes. For example, treatment of wild-type JE2 *S. aureus* with PC190723 (12), an FtsZ inhibitor, induces a distinct phenotype. Cells become enlarged and are arrested in the first stage of the cell cycle (14), a stage typically associated with increased roundness. As illustrated in panel a of Fig. 3, mAIcrobe accurately detects and quantifies these morphological changes, underscoring its utility in phenotypic drug screening.

**Fig. 3.**
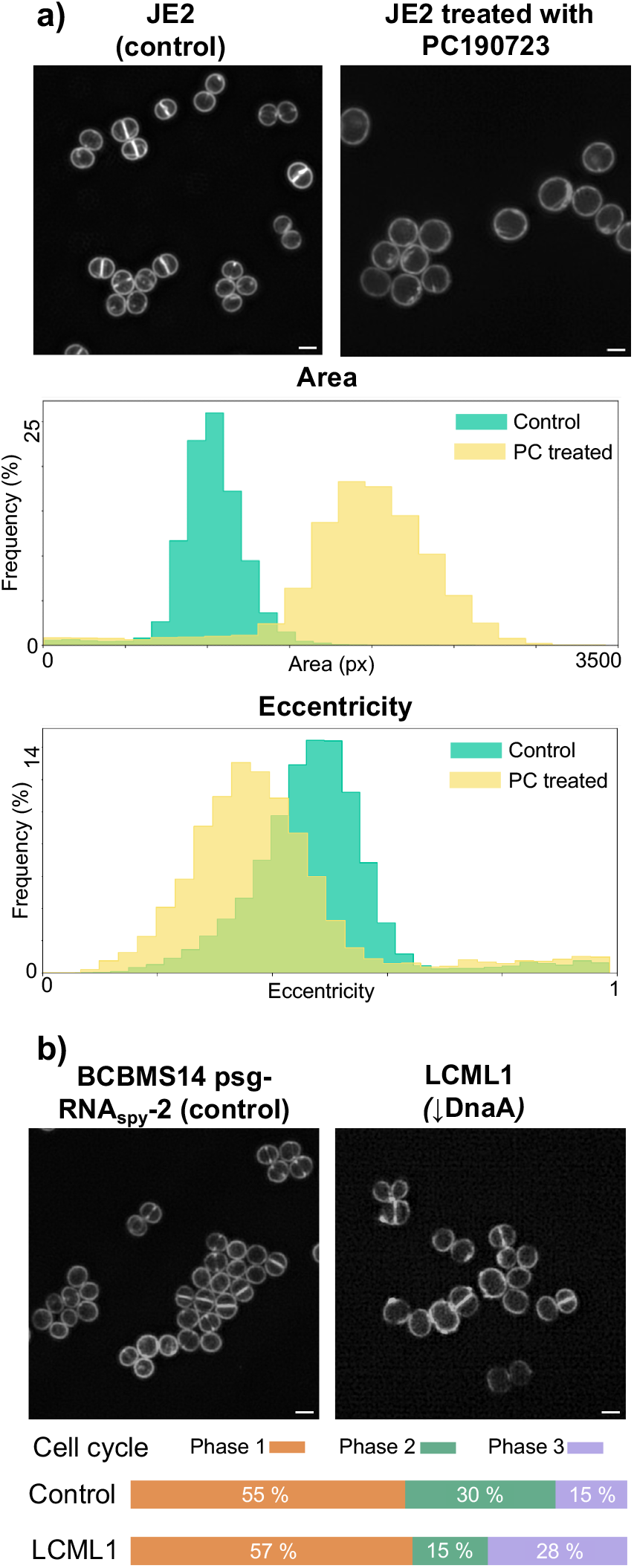
Quantitative phenotyping with mAIcrobe. mAIcrobe is capable of identifying phenotypic variations in microbial cells. a) Morphological changes of *S. aureus* cells treated with the antibiotic PC190723 (12), which leads to larger and rounder cells. The histograms show cell area and eccentricity for control (green, n=12831 cells) and PC190723 treated (yellow, n=4705 cells) JE2 *S. aureus* cells. b) Analysis of cell cycle progression in a *S. aureus* strain (control strain BCBMS14 psg-RNA_spy_-2, n=3429 cells) following CRISPR interference-mediated knockdown of *dnaA* expression (LCML1, n=1647 cells) (13). Cells are classified into three distinct cell cycle phases. Phase 1: round cells with no discernable septa; Phase 2: cells that started to elongate with an open septa; Phase 3: cells with fully closed septa. All scale bars are 1 µm.

### Classification

A principal strength of mAIcrobe is its adaptable classification system, which is powered by a convolutional neural network (CNN). This system is designed for flexibility and can be fine-tuned to address a variety of biological questions, including cell cycle analysis and antibiotic phenotyping.

The classification module employs a CNN architecture previously developed for cell cycle analysis of *S. aureus* (5). For example, on images of *S. aureus* where the essential DNA replication initiator protein DnaA (15, 16) was depleted using CRISPR interference (CRISPRi) (13), mAIcrobe identified altered cell cycle progression (Fig. 3b). This analysis reveals quantifiable differences in cell division timing, which may provide new insights into the role of DnaA in cell cycle regulation.

To support adaptation to diverse experimental conditions and applications beyond cell cycle analysis, a codeless Jupyter notebook (Table S1) is provided for straightforward model retraining and fine-tuning. This approach lowers barriers to the development of custom analysis pipelines. The adaptability of the classification system is demonstrated in Fig. 4, which shows the *S. aureus* cell cycle model retrained for antibiotic phenotype detection in *Escherichia coli* (Fig. S2) (6).

**Fig. 4.**
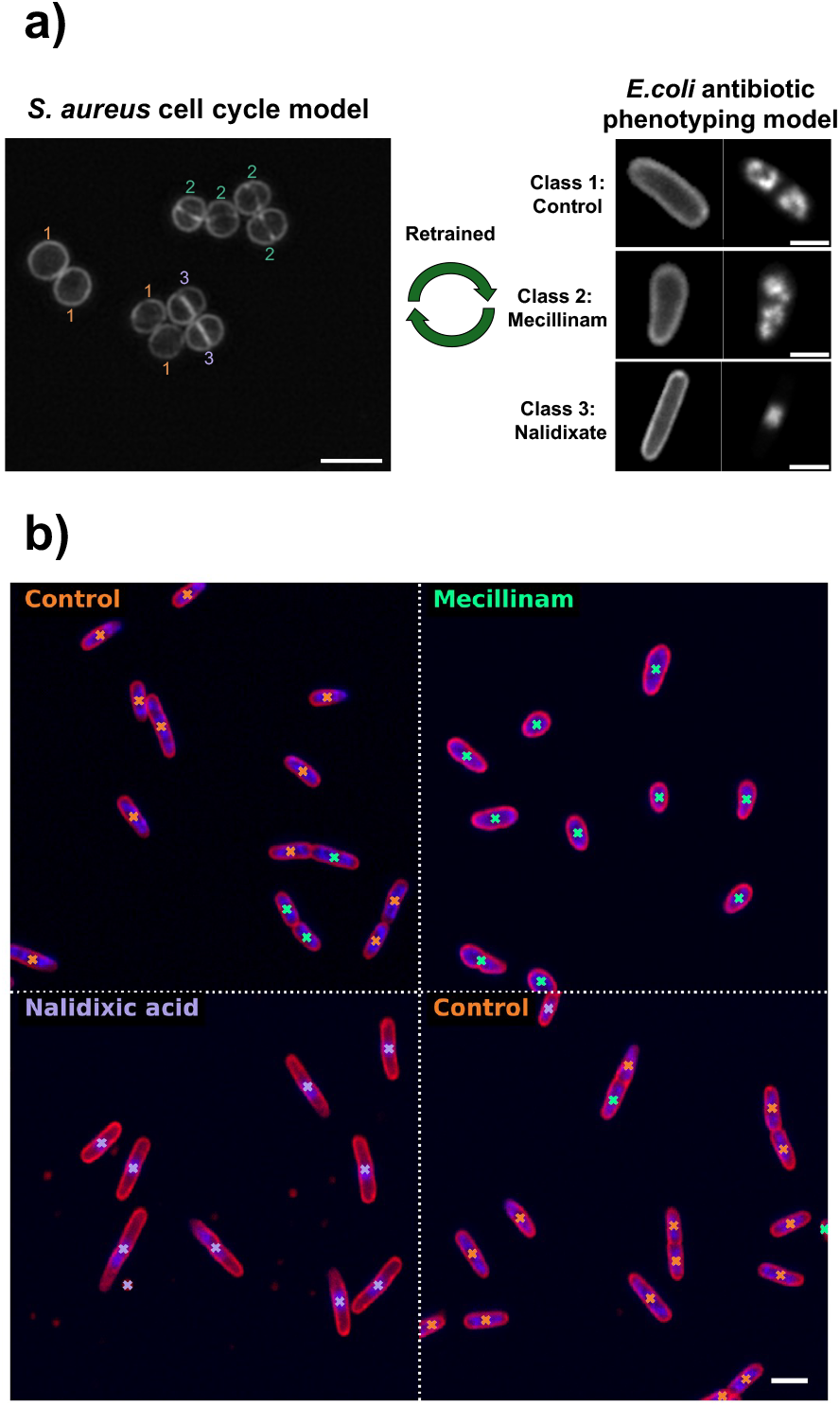
Adaptable classification model in mAIcrobe. a) SIM image of *S. aureus* labeled with membrane dye NileRed. Orange, green, and purple numbers indicate automatically classified cells in phases 1, 2, or 3, respectively, using mAIcrobe’s pretained classification model. b) Synthetic image obtained by stitching together multiple fields of view showcasing different drug treatments. mAIcrobe classification model was fine-tuned to classify *E. coli* cells as control or showcasing the effects of different antibiotics (mecillinam and nalidixic acid). Small crosses indicate classification results (orange for control, green for mecillinam and purple for nalidixic acid). Scale bars are 2 µm (*S. aureus* panel a) and 3 µm (*E. coli* in panel a and b).

## Discussion

mAIcrobe addresses key limitations in current bacterial image analysis workflows by offering a unified framework that integrates deep learning approaches with practical accessibility. Offering a variety of segmentation models constitutes a substantial improvement over single-algorithm methods, as demonstrated by comparative analysis across diverse bacterial morphologies and imaging modalities. Although tools such as eHooke have contributed significantly to the field, they remain constrained by algorithm-specific limitations and morphological restrictions, which reduce their broader applicability.

The integration of StarDist, CellPose, and custom U-Net models within mAIcrobe enables the selection of optimal segmentation approaches for specific experimental conditions. This flexibility is essential given the morphological diversity among bacterial species and the range of microscopy techniques used in contemporary microbiology. Validation across *S. aureus, E. coli, S. pneumoniae*, and *B. subtilis* demonstrates that the multi-model approach maintains high segmentation accuracy while accommodating diverse cell shapes and imaging protocols.

The adaptable classification system constitutes a key innovation, facilitating the transition from fixed-purpose tools to customisable analysis platforms. By offering accessible retraining protocols through Jupyter notebooks, mAIcrobe reduces technical barriers that have previously limited the adoption of machine learning in bacterial microscopy. The adaptation of the model from *S. aureus* cell cycle classification to *E. coli* antibiotic phenotyping demonstrates the framework’s capacity to address diverse biological questions. Jupyter notebooks, which can be used locally or via Google Colab, enable users to retrain the classification model with minimal computational expertise (Table S1).

The morphological measurement capabilities enable comprehensive quantitative profiling of bacterial phenotypes. This functionality is particularly valuable for detecting morphological changes indicative of key biological processes, as demonstrated in analyses of DnaA depletion effects and antibiotic-induced morphological alterations. The ability to export quantitative data in standard formats facilitates integration with statistical analysis workflows and supports reproducible research.

Integration with the napari ecosystem offers strategic advantages for long-term sustainability and community adoption. In contrast to standalone software requiring independent maintenance and feature development, napari plugins benefit from shared infrastructure, advanced visualisation capabilities, and an active development community. This approach ensures that mAIcrobe evolves in parallel with advances in the broader image analysis field while maintaining compatibility with complementary tools.

## Conclusions

mAIcrobe offers a comprehensive set of computational tools for bacterial microscopy analysis, delivering a unified solution to the fragmented landscape of existing software. The principal innovation of the framework is its seamless integration of multiple segmentation algorithms with adaptable classification models, enabling comprehensive analysis across diverse bacterial species and experimental conditions without requiring transitions between different software packages.

Empirical validation indicates that mAIcrobe’s multi-model approach maintains high analytical performance while substantially expanding the range of addressable biological questions. Demonstrated applications, including the detection of cell cycle defects in DnaA-depleted *S. aureus* and the characterisation of antibiotic-induced morphological changes, highlight the framework’s capacity to reveal biologically relevant phenotypes.

Integration with the napari ecosystem positions mAIcrobe as a forward-looking solution that addresses both current analytical needs and future scalability requirements. By leveraging napari’s extensible architecture and active development community, the framework ensures long-term sustainability and seamless integration with complementary analysis tools. The open-source implementation and accessible retraining protocols broaden access to advanced image analysis capabilities,

potentially accelerating discovery across multiple areas of bacterial cell biology.

With the growing demand for sophisticated analytical approaches to address complex biological questions, mAIcrobe provides a robust foundation for next-generation bacterial microscopy analysis. The modular design and extensible architecture enable the incorporation of future methodological advances while maintaining the accessibility and reliability necessary for routine research. This combination of current capability and future adaptability establishes mAIcrobe as a valuable addition to the computational microbiology toolkit.

## ABOUT THIS MANUSCRIPT

This manuscript was prepared using R*χ*iv-Maker v1.22.1 (17). This work is licensed under CC BY 4.0.

## DATA AVAILABILITY

All the segmentation models and the classification model are available via the mAIcrobe GitHub repository (https://github.com/henriqueslab/maicrobe) All the data used in this study is publicly available. The datasets of *B. subtilis* and *E. coli* are available in (6). The datasets of *S. aureus* strains, JE2, BCBMS14 psg-RNA_spy_-2 and LCML1, and the dataset of Pen6 *S. pneumoniae* were acquired in-house and are available on Zenodo (https://doi.org/10.5281/zenodo.17306839).

A comprehensive list of all models and the respective training and test datasets can be found in Table S5.

## CODE AVAILABILITY

Source code for the napari-mAIcrobe plugin can be accessed via the GitHub repository (https://github.com/henriqueslab/maicrobe). The plugin is also available and installable through PyPi (https://pypi.org/project/napari-mAIcrobe/). Notebooks for training the classification model can be found in the notebooks folder of the mAIcrobe repository.

## AUTHOR CONTRIBUTIONS

A.D.B., M.G.P., B.M.S. and R.H. designed the study. A.D.B. developed the code and trained the models. A.D.B., B.M.S. and D.A. prepared the samples and acquired the *S. aureus* data. B.P.M. and S.R.F. prepared the *S. pneumoniae* samples and A.D.B acquired the data. B.M.S., M.G.P. and R.H. supervised the project. A.D.B., M.G.P., B.M.S. and R.H. wrote the manuscript with input from all authors.

## ACKNOWLEDGEMENTS

A.D.B. acknowledges the FCT 2021.06849.BD fellowship. D.A. acknowledges the FCT 2022.12215.BD fellowship. B.P.M. acknowledges the FCT UI/BD/151527/2021 fellowship.

## EXTENDED AUTHOR INFORMATION

- **António D. Brito:**
- **Dominik Alwardt:**
- **Beatriz de P. Mariz:**
- **Sérgio R. Filipe:**
- **Mariana G. Pinho:**
- **Bruno M. Saraiva:**
[in bsaraiva](https://linkedin.com/in/bsaraiva)
- **Ricardo Henriques:**
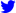 [HenriquesLab](https://x.com/HenriquesLab); * henriqueslab.bsky.social;

## Methods

### Image acquisition

The datasets of *S. aureus* strains, JE2 (15), BCBMS14 psg-RNA_spy_-2 and LCML1 (13) (Table S4) were acquired in-house.

For the growth of BCBMS14 psg-RNA_spy_-2 and LCML1 overnight cultures of both strains were back-diluted 1:500 into 10 mL of fresh tryptic soy broth (TSB, Difco) media containing 10 µg/ml chloramphenicol (Sigma-Aldrich) and grown at 37 °C for 1 hour. After 1 hour, anhydrotetracycline (aTc, Sigma-Aldrich) was added to the medium to a final concentration of 100 ng/ml. After another hour, a 1 mL aliquot of each culture was incubated with 2.5 µg/mL NileRed (Invitrogen) for 5 min at 37 °C with shaking. The culture was pelleted (10000 rpm for 1 min), supernatant was removed and the pellet was resuspended in 30 µL of phosphate-buffered saline (PBS, NaCl 137 mM, KCl 2.7 mM, Na_2_HPO_4_10 mM, KH_2_PO_4_ 1.8 mM). One microliter of the resuspended culture was then placed on a thin layer of 1.2% (w/v) agarose (TopVision Thermo Fisher Scientific) in PBS and imaged via structured illumination microscopy (SIM).

For the growth of untreated JE2, an overnight culture was back-diluted 1:200 into 10 mL of fresh TSB media and grown at 37 °C until cells reached mid-exponential growth phase (OD600 of 0.8). Afterwards, a 1 mL aliquot of culture was incubated with 5 µg/mL NileRed (Invitrogen) and 1 µg/mL Hoechst 33342 (Invitrogen) for 5 min at 37 °C with shaking. Culture was then centrifuged, washed with 1 ml of 1:3 (vol/vol) TSB/PBS solution, and resuspended in 20 µL of the same solution. Cells were mounted on microscope slides covered with a layer of 1.2% (w/v) agarose in PBS and imaged via structured illumination microscopy (SIM).

SIM was performed using an Elyra PS.1 microscope (Zeiss) with a Plan-Apochromat 63x/1.4 oil DIC M27 objective. SIM images were acquired using three grid rotations, with a 34-µm grating period for the 561-nm laser (100 mW) and 23 µm grating period for the 405-nm laser (50 mW). Images were captured using a Pco.edge 5.5 camera and reconstructed using ZEN software (black edition, 2012; version 8.1.0.484) on the basis of a structured illumination algorithm, with synthetic, channel-specific optical transfer functions and noise filter settings ranging from 6 to 8.

The dataset of Pen6 *S. pneumoniae* strains was also acquired in-house. Briefly, overnight cultures were back-diluted 1:50 into 5 mL of fresh C medium supplemented with yeast extract (0.8% Difco Laboratories) (C+Y media). C medium was prepared as described in (18). Cells were grown at 37 °C to early exponential phase (OD600 0.2-0.3). A 1 mL aliquot of the culture was centrifuged (10000 rpm for 1 min) and the pellet was resuspended in 30 µL of pre-C medium (18). Two microliters of the resuspended culture were then placed on a thin layer of 1.2% (w/v) agarose in pre-C (18) media and imaged using a Zeiss Axio Observer microscope equipped with a Plan-Apochromat 100x/1.4 oil Ph3 objective, a Retiga R1 CCD camera (QImaging), a white-light source HXP 120 V (Zeiss) and the software ZEN blue v2.0.0.0 (Zeiss).

### Biological image datasets

Datasets of *B. subtilis* expressing FtsZ-GFP (strain SH130, PY79 Δ*hag ftsZ::ftsZ-gfp-cam* (19)), which was used to train a U-Net segmentation model, and *E. coli* (strain NO34 (20)) exposed to various antibiotics, which was used to train a classification network, are publicly available in (6) alongside their annotations.

The dataset of Pen6 *S. pneumoniae* that was used to test and train a U-Net segmentation model, was acquired in-house and is available on Zenodo (https://doi.org/10.5281/zenodo.17306839).

The *S. aureus* dataset containing untreated and PC190723 treated JE2 cells labeled with NileRed is publicly available in (21). The same dataset alongside in-house acquired images of BCBMS14 psg-RNA_spy_-2 was used to train and test a StarDist segmentation model and can be found in Zenodo (https://doi.org/10.5281/zenodo.17306839).

The dataset WT JE2 *S. aureus* cells labeled with NileRed and Hoechst, used for validating morphometrics, is available in Zenodo (https://doi.org/10.5281/zenodo.17306839).

The *S. aureus* dataset containing CRISPRi-depleted strain (LCML1) and its respective control (BCBMS14 psg-RNA_spy_-2) was used to test the pretrained *S. aureus* cell cycle classification model. The dataset was acquired in-house and is available on Zenodo (https://doi.org/10.5281/zenodo.17306839).

A comprehensive list of all biological datasets used in this study can be found in Table S5.

### Segmentation networks

The StarDist model used for *S. aureus* segmentation was trained using a dataset of untreated (10 FoVs) and PC190723 treated (12 FoVs) JE2 *S. aureus* labeled with NileRed.The training dataset is the dataset available in (21). The test dataset contains 3 FoVs of BCBMS14 psg-RNA_spy_-2 *S. aureus* strain labeled with NileRed (Table S4) (13). Both the training and the test dataset are deposited in Zenodo (https://doi.org/10.5281/zenodo.17306839). Training was performed on a Jupyter notebook, adapted from the example notebooks provided by StarDist authors, that can be found in the code repository of this work (Table S1).

The U-Net model used for *S. pneumoniae* and *B. subtilis* segmentation was trained on an adapted ZeroCostDL4Mic 2D U-Net notebook (7), that can be found in the code repository of this work (Table S1). The U-Net model was trained to identify background, cell edge, and cell interior. To obtain the final label image, scikit-image’s (22) watershed segmentation was used (23). First, a mask image is generated by performing the binary union of the cell edge and cell interior. The input to the watershed algorithm is the inverted mask alongside the cell interiors as marker basins. The training and test datasets of *S. pneumoniae* were obtained in-house and are available on Zenodo (https://doi.org/10.5281/zenodo.17306839). The *B. subtilis* training and test datasets are publicly available in (6).

For both training datasets, data augmentation was performed using image rotations and flips. The hyperparameters of each model can be found in Table S6.

### Classification network

The classification network trained on *E. coli* data is a convolutional neural network with an architecture described in (5). This network was retrained using the *E. coli* antibiotic phenotyping dataset from (6). The fields of view pertaining to the control condition plus those corresponding to exposure to mecillinam and nalidixic acid were split into the DNA and membrane channels, and the membrane channel was segmented using the CellPose cyto3 model (10). Individual cell crops were extracted from the segmented fields of view using mAIcrobe to generate the final dataset needed for training and testing. In total, the training dataset contained 1164 *E. coli* cell crops while the test dataset contained 416 cells. The network was trained for 200 epochs with a batch size of 32 and a learning rate of 0.001 and a validation split of 20%. Data augmentation was performed using Keras (24) RandomRotation and RandomFlip preprocessing layers. These layers, at training time only, perform random horizontal and vertical flipping and random rotations between -180º and 180º. Training was done using a Jupyter notebook (25) available in our GitHub repository (Table S1).

## Supplementary Information

**sup. Table S1.**
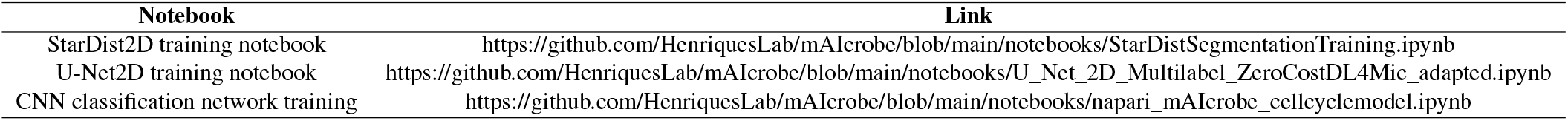
Jupyter notebooks available as part of mAIcrobe.

**sup. Table S2.** Experimental conditions for datasets used in this study.

| Description | Species | Microscopy modality | Reference |
| --- | --- | --- | --- |
| *S. aureus* datasets | *S. aureus* | SIM | This study, (21) |
| *S. pneumoniae* U-Net dataset | *S. pneumoniae* | Widefield (phase contrast) | This study |
| *B. subtilis* U-Net dataset | *B. subtilis* | Widefield (fluorescence) | (6) |
| *E. coli* classification model dataset | *E. coli* | Widefield (fluorescence) | (6) |

**sup. Table S3.** Performance metrics for the segmentation networks used in this study. Average Intersection over Union (IoU), Recall and Precision values were calculated on the respective test datasets. All tests datasets had three FoV’s.

| Model | Average IoU | Recall | Precision |
| --- | --- | --- | --- |
| StarDist *S.aureus* | 0.999 | 0.994 | 0.872 |
| U-Net *S.pneumoniae* | 0.998 | 0.988 | 0.898 |
| U-Net *B.subtilis* | 0.923 | 1.00 | 0.840 |

**sup. Table S4.**
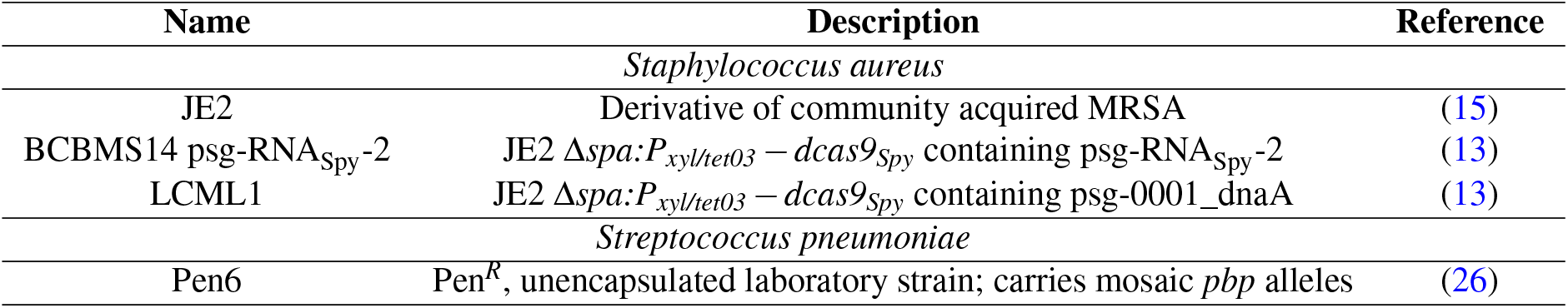
Strains used in this study.

**sup. Table S5.**
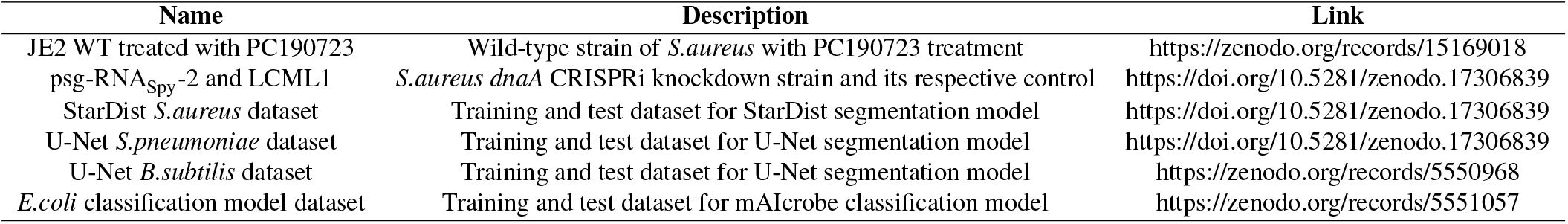
Datasets used in this study and their respective repository links.

**sup. Table S6.** Hyperparameters for the segmentation networks used in this study.

| Organism | Microscopy | Network | Train/test images | Epochs | Steps | Image size | Patch size | Batch size | Learning rate | %Validation | Augmentation | Misc |
| --- | --- | --- | --- | --- | --- | --- | --- | --- | --- | --- | --- | --- |
| *S.aureus* | SIM | StarDist | 20/3 | 400 | 100 | 2430x2430 | 256x256 | 4 | 0.0003 | 15 | Random flip and intensity changes | grid 2, 32 rays |
| *S.pneumoniae* | Phase Contrast | U-Net | 16/3 | 100 | 70 | 1392x1040 | 256x256 | 4 | 0.0003 | 10 | VHFlip and 180 | Finetuned from a *S.aureus* U-Net model |
| *B.subtilis* | Fluorescence | U-Net | 7/3 | 100 | 6 | 1024x1024 | 512x512 | 4 | 0.0005 | 10 | VHFlip and 180°rot | Finetuned from a *S.aureus* U-Net model |

**sup. Fig. S1.**
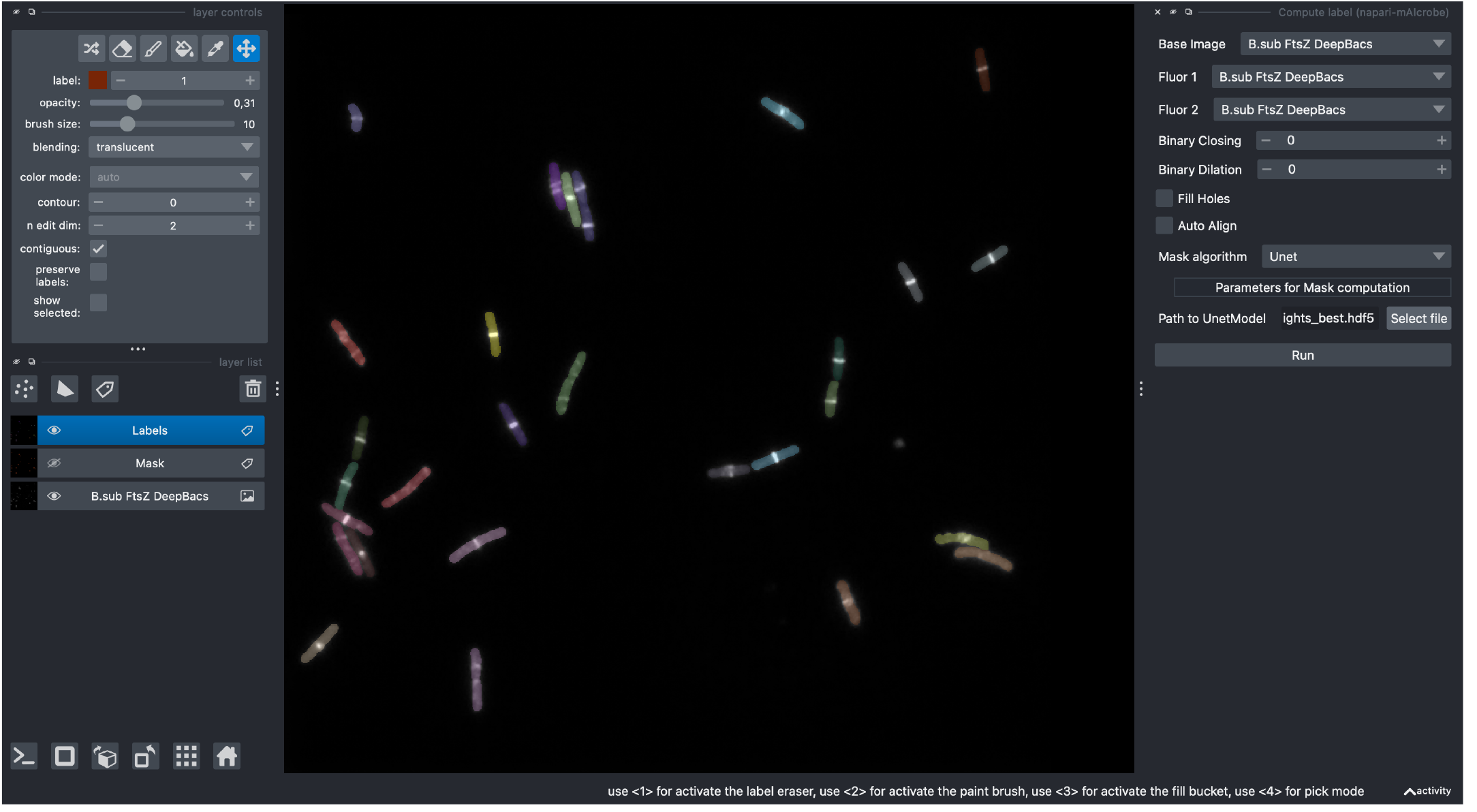
Example screenshot of the mAIcrobe napari plugin. In this screenshot, we showcase an image of *B*.*subtilis* cells being segmented using a U-Net model loaded into the mAIcrobe plugin. The plugin’s segmentation interface is visible on the right side of the image, displaying various options and settings for segmentation, that dynamically change according to the segmentation model chosen.

**sup. Fig. S2.**
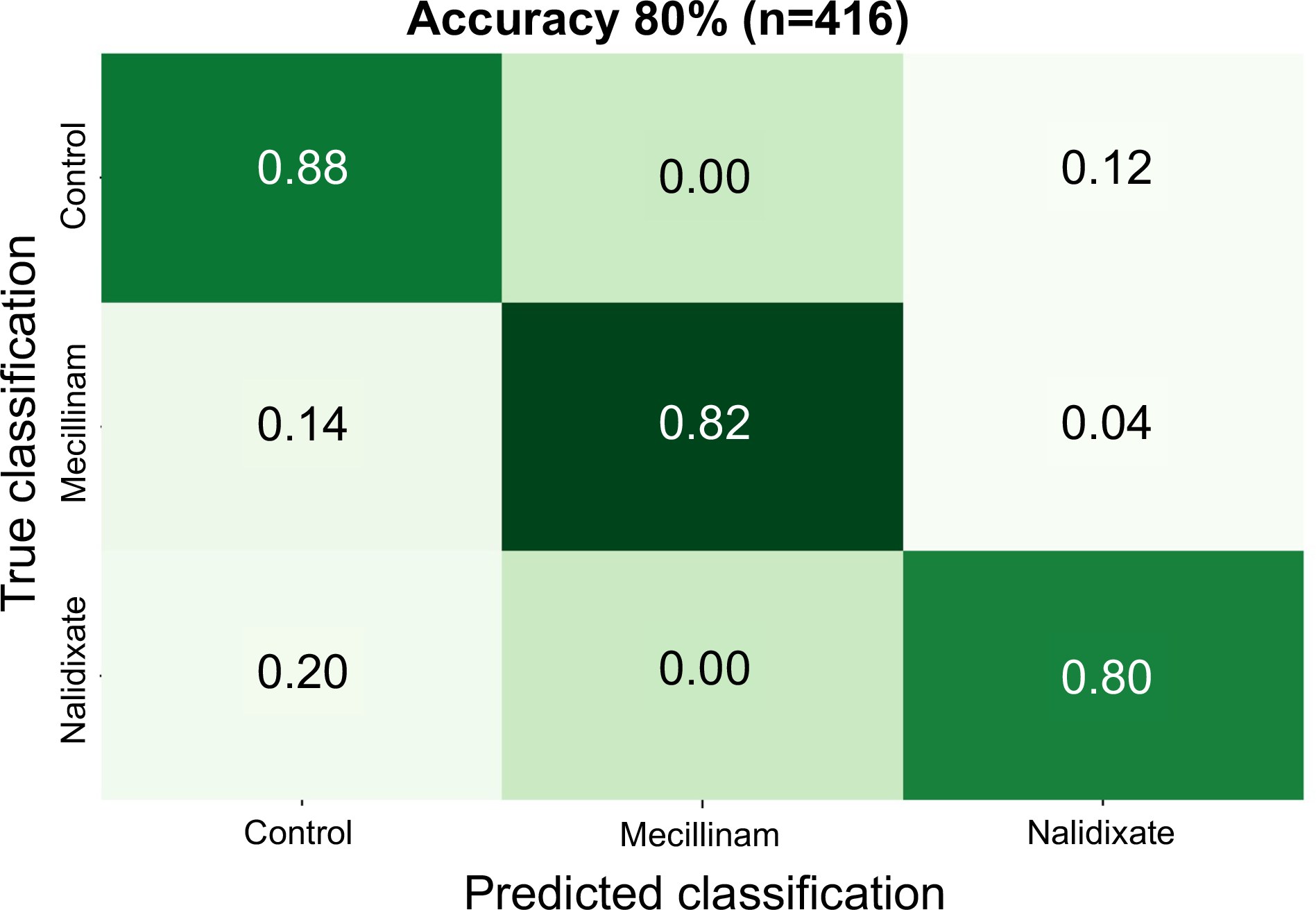
Confusion matrix for E. coli classification model. The confusion matrix shows the performance of the retrained mAIcrobe CNN on a test dataset of 416 cells.

